# CellPhoneDB v2.0: Inferring cell-cell communication from combined expression of multi-subunit receptor-ligand complexes

**DOI:** 10.1101/680926

**Authors:** Mirjana Efremova, Miquel Vento-Tormo, Sarah A. Teichmann, Roser Vento-Tormo

**Affiliations:** Wellcome Sanger Institute, Wellcome Genome Campus, Cambridge CB10 1SD, UK; YDEVS software development, Valencia, 46009, Spain; Theory of Condensed Matter Group, Cavendish Laboratory, University of Cambridge, JJ Thomson Ave, Cambridge CB3 0EH, UK

## Abstract

Cell-cell communication mediated by receptor-ligand complexes is crucial for coordinating diverse biological processes, such as development, differentiation and responses to infection. In order to understand how the context-dependent crosstalk of different cell types enables physiological processes to proceed, we developed CellPhoneDB, a novel repository of ligands, receptors and their interactions^1^. Our repository takes into account the subunit architecture of both ligands and receptors, representing heteromeric complexes accurately. We integrated our resource with a statistical framework that predicts enriched cellular interactions between two cell types from single-cell transcriptomics data. Here, we outline the structure and content of our repository, the procedures for inferring cell-cell communication networks from scRNA-seq data and present a practical step-by-step guide to help implement the protocol. CellPhoneDB v2.0 is a novel version of our resource that incorporates additional functionalities to allow users to introduce new interacting molecules and reduce the time and resources needed to interrogate large datasets. CellPhoneDB v2.0 is publicly available at https://github.com/Teichlab/cellphonedb and as a user-friendly web interface at http://www.cellphonedb.org/. In our protocol, we demonstrate how to reveal meaningful biological discoveries from CellPhoneDB v2.0 using published data sets.

## Introduction

We developed CellPhoneDB, a public repository of ligands, receptors and their interactions to enable a comprehensive, systematic analysis of cell–cell communication molecules. Our repository relies on the use of public resources to annotate receptors and ligands as well as manual curation of specific families of proteins involved in cell-cell communication. We include subunit architecture for both ligands and receptors to represent heteromeric complexes accurately (Figure 1). This is crucial, as cell-cell communication relies on multi-subunit protein complexes that go beyond the binary representation used in most databases and studies. In order to integrate all the information in a flexible, distributable and amendable environment, we developed an SQLite relational database. CellPhoneDB v2.0 enables systematic updates of the stored information in database releases, as well as the continuous submission of novel interacting partners by users.

**Figure 1.**
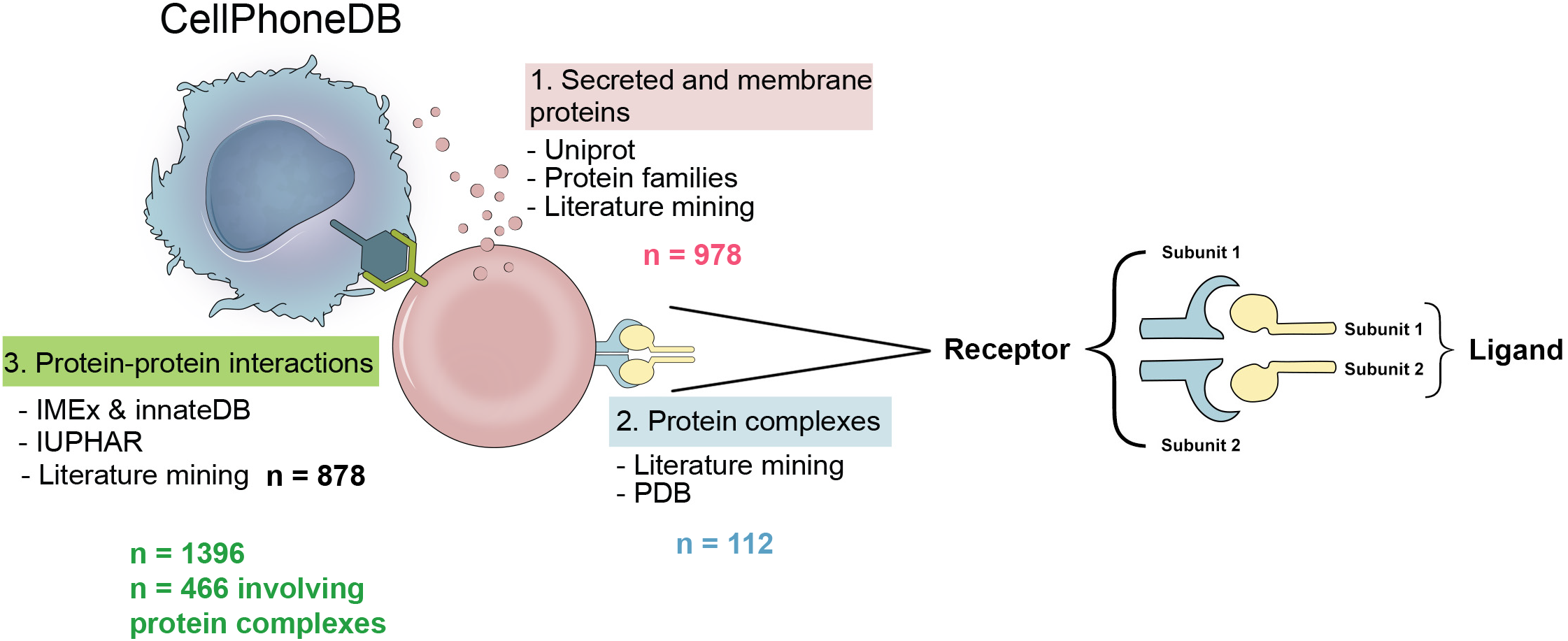
Overview of the database. **a**, Information aggregated within www.CellPhoneDB.org. CellPhoneDB stores a total of 978 proteins, 501 are secreted proteins and 585 are membrane proteins. These proteins are involved in 1396 interactions, out of those 466 are heteromers. There are 474 interactions that involve secreted proteins and 490 interactions that involve only membrane proteins. There are a total of 250 interactions that involve integrins.

Our repository forms the basis of a computational approach to identify biologically relevant interacting ligand-receptor partners from single-cell transcriptomics (scRNAseq) data. We consider the expression levels of ligands and receptors within each cell type, and use empirical shuffling to calculate which ligand–receptor pairs display significant cell-type specificity (Figure 2). This predicts molecular interactions between cell populations via specific protein complexes and generates potential cell–cell communication networks. Specificity of the ligand-receptor interaction is important, as some of the ligand-receptor pairs are ubiquitously expressed by the cells in a tissue, and therefore not informative regarding specific communication between particular cell states. The computational code is available in github https://github.com/Teichlab/cellphonedb. In addition, we developed a user-friendly web interface at www.CellPhoneDB.org, where users can search for ligand–receptor complexes, interrogate their own scRNAseq data and download and visualise their results. Our new database implementation now offers improved visualization capability including dot plots and heatmaps.

**Figure 2.**
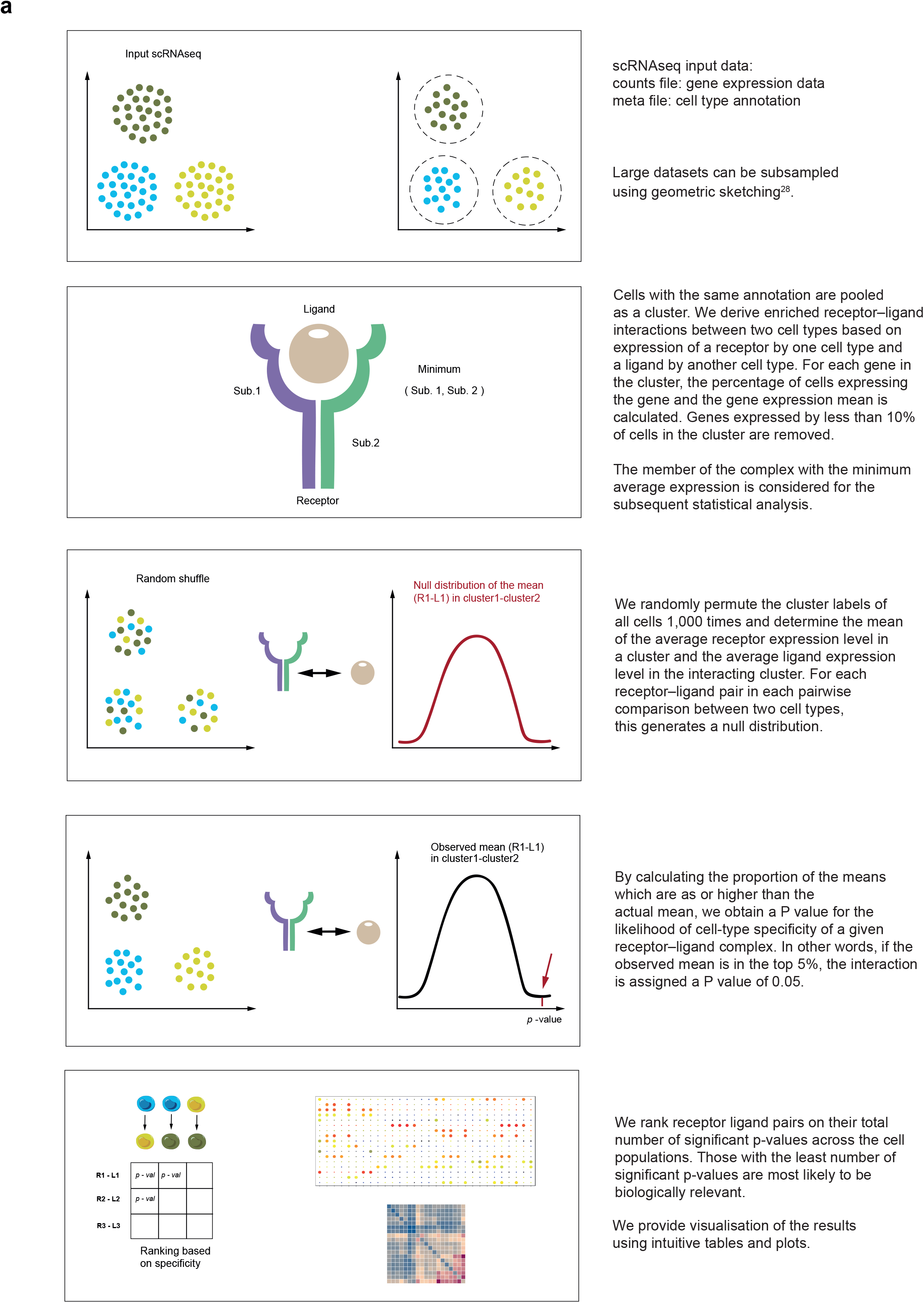
Overview of the statistical method. **a**, Statistical framework used to infer ligand– receptor complex specific to two cell types from single-cell transcriptomics data. Predicted P values for a ligand–receptor complex across two cell clusters are calculated using permutations, in which cells are randomly re-assigned to cluster.

We originally applied this computational framework to study maternal-fetal communication at the decidual-placental interface during early pregnancy^1^. Briefly, our analysis revealed new immunoregulatory mechanisms and cytokine signaling networks existing between the cells in the maternal-fetal interface, which guarantee the coexistence of both the mother and developing fetus (Figure 3). In the present protocol, we describe and discuss in detail how this analysis can be carried out, using our maternal-fetal study as an illustration.

**Figure 3.**
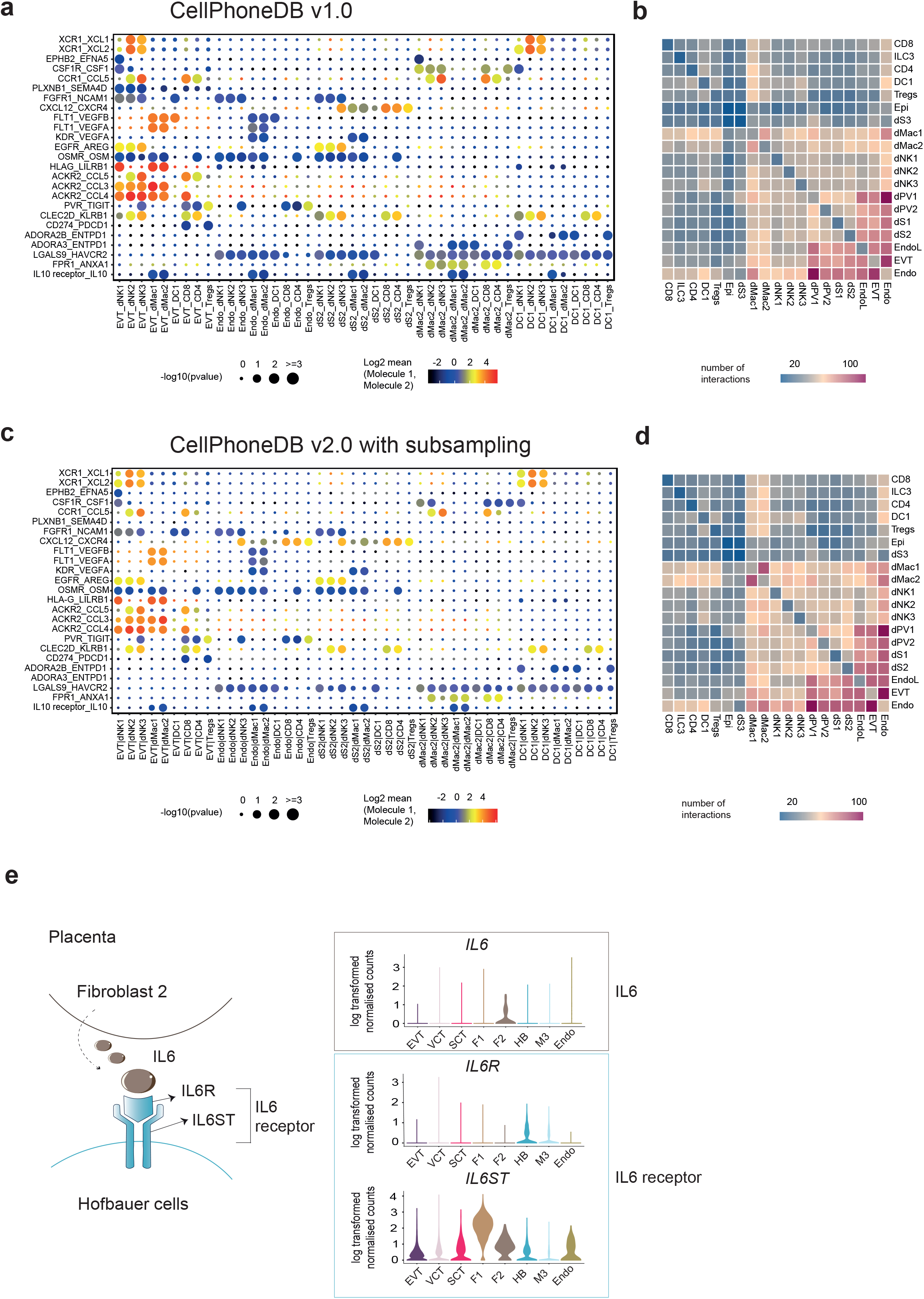
Diagram generation lists. **a**, Basic steps in the generation of lists to populate the tables in CellPhoneDB.

The protocol is generalizable to any other dataset containing potentially interacting cell populations. For example, CellPhoneDB helped us identify a shift in the cellular communication from a network that was dominated by mesenchymal-epithelial interactions in healthy airways, to a Th2 cell--dominated interactome in asthmatic airways^2^. In another recent example, our cell-cell communication framework revealed the complex interplay among diverse cells in the evolving tumor microenvironment of a murine melanoma model where multiple immunosuppressive mechanisms coexist within a heterogeneous stromal compartment^3^. We anticipate that the information stored in CellPhoneDB 2.0 will have the potential to resolve cellular communication when combined with the spatial location of the cells that can be quantified using highly multiplexed spatial methods ^4–7^.

Single cell transcriptomics technologies have provided a unique opportunity to analyse the expression of multiple cell types and systematically decode intercellular communication networks. There are now several other published methods to infer potentially relevant interactions between two cell populations from scRNAseq. The majority of these methods use lists of binary ligand-receptor pairs to assign communication between cells, without considering multimeric receptors. Relevant interactions are inferred by filtering based on the expression level of the ligand and receptor. In these methods, only the interaction pairs that pass a certain threshold of cells expressing the specific interactors in the respective cell populations are selected for the downstream analysis ^8–13^. For example, in addition to filtering based on expression level, Cohen et al.^14^ used hierarchical clustering with Spearman correlation to identify ligand/receptor modules and construct an interaction graph. Others, such as Kumar et al^15^, scored interactions by calculating the product of average receptor and average ligand expression in the corresponding cell types and used a one-sided Wilcoxon rank-sum test to assess the statistical significance of each interaction score. Halpern et al.^16^ computed a z-score of the mean of each interacting molecule in each cluster to calculate the enrichment of each ligand and receptor in each cluster. To test for enrichment of the number of receptor-ligand pairs between two cell populations, Joost et al.^17^ performed random sampling of receptors and ligands and compared this number with the observed number of receptor-ligand pairs. In a similar way, Boisset et al.^17, 18^ applied cluster label permutations to create a distribution of the number of chance interacting pairs between cell populations and then compared this to the actual number of interactions to identify enriched or depleted interactions compared with the numbers in the background model.

A major strength of CellPhoneDB beyond most other databases is that it takes into account the structural composition of ligands and receptors, which often involve the interaction of multiple subunits. This is particularly clear for protein families like many of the cytokine families, where receptors share structural subunits, and the affinity of the ligand is determined by the specific combination of the subunits (Figure 3e). Roughly one third of the receptor-ligand complexes in our database have a multi-subunit stoichiometry greater than binary one-to-one interactions. Specifically, there are 466 interactions in our repository which involve heteromers, and 163 of them comprise cytokines.

CellPhoneDB v2.0 has new features, such as subsampling of the original dataset to enable the fast query of large datasets, and visualisation of the results using intuitive tables and plots. The statistical analysis package of CellPhoneDB is available as a user-friendly web interface (CellPhoneDB.org) and an easy-to-use python package available on github.

## Database design and analysis method

### Database input files

CellPhoneDB stores ligand-receptor interactions as well as other properties of the interacting partners, including their subunit architecture and gene and protein identifiers. In order to create the content of the database, four main .csv data files are required: “genes_input.csv”, “proteins_input.csv”, “ complexes_input.csv” and “interactions_input.csv” (Figure 4).

**Figure 4.**
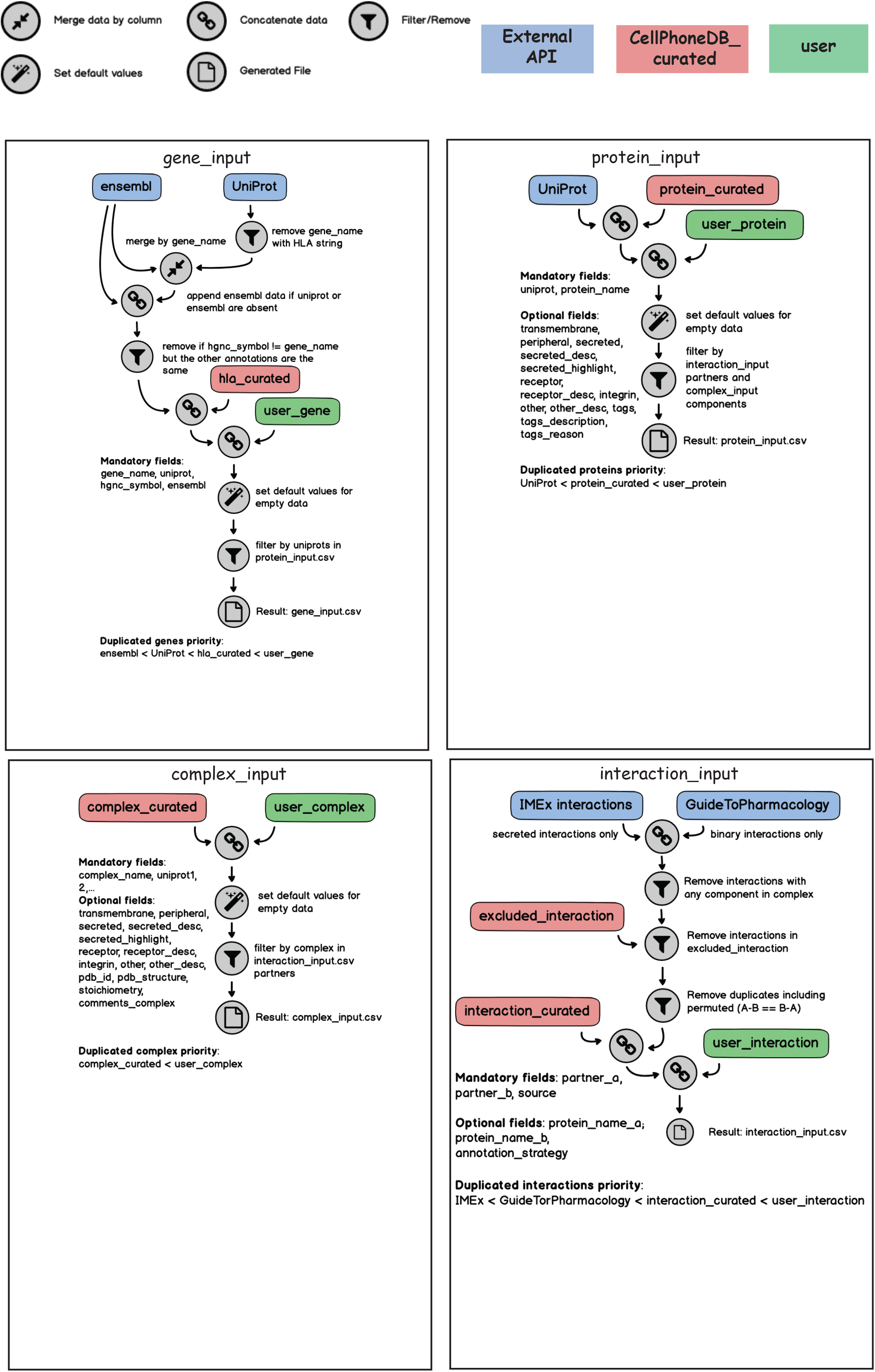
Example dataset run with CellPhoneDB and CellPhoneDB v2.0. **a**, Overview of selected ligand–receptor interactions using CellPhoneDB on the decidua dataset from ^1^; P values indicated by circle size, scale on right. The means of the average expression level of interacting molecule 1 in cluster 1 and interacting molecule 2 in cluster 2 are indicated by colour. **b**, Heatmap showing the total number of interactions between cell types in the decidua dataset obtained with CellPhoneDB. **c**, Overview of selected ligand–receptor interactions using the CellPhoneDB v2.0 with subsampling on the decidua dataset. P values indicated by circle size, scale on right. The means of the average expression level of interacting molecule 1 in cluster 1 and interacting molecule 2 in cluster 2 are indicated by colour. ⅓ of the dataset was subsampled. **d**, Heatmap showing the total number of interactions between cell types in the decidua dataset obtained with CellPhoneDB v2.0 with subsampling. ⅓ of the dataset was subsampled. **e**. An example of significant interactions involving complexes identified by CellPhoneDB in the placenta dataset ^1^. Violin plots show log-transformed, normalized expression levels of the components of the IL6–IL6R complex in placental cells. IL6 expression is enriched in the fibroblast 2 cluster and the two subunits of the IL6 receptors (IL6R and IL6ST) are co-expressed in Hofbauer cells.

#### 1. “gene_input”

*Mandatory fields: “gene_name”; “uniprot”; “hgnc_symbol” and “ensembl”*

This file is crucial for establishing the link between the scRNAseq data and the interaction pairs stored at the protein level. It includes the following gene and protein identifiers: i) gene name (“gene_name”); ii) UniProt identifier (“uniprot”); iiii) HUGO nomenclature committee symbol (HGNC) (“hgnc_symbol”) and iv) gene ensembl identifier (ENSG) (“ensembl”). In order to create this file, lists of linked proteins and genes identifiers are downloaded from UniProt and merged using gene names. Several rules need to be considered when merging the files:

- UniProt annotation prevails over the gene Ensembl annotation when the same gene Ensembl points towards different UniProt identifiers.
- UniProt and Ensembl lists are also merged by their UniProt identifier but this information is only used when the UniProt or Ensembl identifier are missing in the original merged list by gene name.
- If the same gene name points towards different HGNC symbols, only the HGNC symbol matching the gene name annotation is considered.
- Only one HLA isoform is considered in our interaction analysis and it is stored in a manually HLA-curated list of genes, named “HLA_curated”.

#### 2. “protein_input”

*Mandatory fields: “uniprot”; “protein_name” Optional fields: “transmembrane”; “peripheral”; “secreted”; “secreted_desc”; “secreted_highlight”; “receptor”; “receptor_desc”; “integrin”; “other”; “other_desc”; “tags”; “tags_ description”; “tags_reason”*

Two types of input are needed to create this file: i) systematic input using UniProt annotation, and ii) manual input using curated annotation both from developers of CellPhoneDB (“proteins_curated”) and users. For the systematic input, the UniProt identifier (“uniprot”) and the name of the protein (“protein_name”) are downloaded from UniProt. For the curated input, developers and users can introduce additional fields relevant to the future systematic assignment of ligand-receptor interactions (see below the “Systematic input from other databases” section for interaction_list). Importantly, the curated information always has priority over the systematic information.

The optional input is organised using the fields described below:

##### a) Location of the protein in the cell

There are four non-exclusive options: transmembrane (“transmembrane”), peripheral (“peripheral”) and secreted (“secreted”, “secreted_desc” and “secreted_highlight”).

Plasma membrane proteins are downloaded from UniProt using the keyword KW-1003 (cell membrane). Peripheral proteins from the plasma membrane are annotated using the UniProt keyword SL-9903, and the remaining proteins are annotated as transmembrane proteins. We complete our lists of plasma transmembrane proteins by doing an extensive manual curation using literature mining and UniProt description of proteins with transmembrane and immunoglobulin-like domains.

Secreted proteins are downloaded from UniProt using the keyword KW-0964 (secreted), and are further annotated as cytokines (KW-0202), hormones (KW-0372), growth factors (KW-0339) and immune-related using UniProt keywords and manual annotation. Cytokines, hormones, growth factors and other immune-related proteins are indicated as “secreted_highlight” in the protein_input lists. “secreted_desc” indicates a quality for the secreted protein.

All the manually annotated information is carefully tagged and can be identified. Please see the “curation tags” section below.

##### b) Receptors and integrins

Three fields are allocated to annotate receptors or integrins: “receptor”, “receptor_desc” and “integrin”.

Receptors are defined by the UniProt keyword KW-0675. The receptors list is extensively reviewed and new receptors are added based on UniProt description and bibliography revision. Receptors involved in immune-cell communication are carefully curated. For some of the receptors, a short description is included in “receptor_desc”.

Integrin is a manual curation field that indicates the protein is part of the integrin family. All the annotated information is carefully tagged and can be identified. For details, see “curation tags” section below.

##### c) Membrane and secreted proteins not considered for the cell-cell communication analysis (“others”)

We created another column named “others” that consists of proteins that are excluded from our analysis. We also added “others_desc” to add a brief description of the excluded protein. Those proteins include: (i) co-receptors; (ii) nerve-specific receptors such as those related to ear-binding, olfactory receptors, taste receptors and salivary receptors; (iii) small molecule receptors; (iv) immunoglobulin chains; (v) viral and retroviral proteins, pseudogenes, cancer antigens and photoreceptors.

##### d) Curation “tags”

Three fields indicate whether the protein has been manually curated: “tags”, “tags_ description” and “tags_reason”.

The “tags” field is related to the manual curation of a protein and contains three fields: (i) ‘N/A’, which indicates that the protein matched with UniProt description in all fields; (ii) ‘To_add’, which indicates that secreted and/or plasma membrane protein annotation has been added; and (iii) ‘To_comment’, which indicates that the protein is either secreted (KW-0964) or membrane-associated (KW-1003), but that we manually added a specific property of the protein (that is, the protein is annotated as a receptor).

The “tags_reason” field is related to the protein properties and has five possible values: (i) ‘extracellular_add’, which indicates that the protein is manually annotated as plasma membrane; (ii) ‘peripheral_add’, which indicates that the protein is manually annotated as a peripheral protein instead of plasma membrane; (iii) ‘secreted_add’, which indicates that the protein is manually annotated as secreted; (iv) ‘secreted_high’, which indicates that the protein is manually annotated as secreted highlight. For cytokines, hormones, growth factors and other immune-related proteins, the option (v) ‘receptor_add’ indicates that the protein is manually annotated as a receptor.

Finally, the “tags_description” field is a brief description of the protein, function or property related to the manually curated protein.

#### 3. “complex_input”

*Mandatory fields: “complex_name”; “uniprot1, 2,...”*

*Optional fields: “transmembrane”; “peripheral”; “secreted”; “secreted_desc”; “secreted_highlight”; “receptor”; “receptor_desc”; “integrin”; “other”; “other_desc”; “pdb_id”; “pdb_structure”; “stoichiometry”; “comments_complex”*

Heteromeric receptors and ligands - that is, proteins that are complexes of multiple gene products - are annotated by reviewing the literature and UniProt descriptions. Cytokine complexes, TGF family complexes and integrin complexes are carefully annotated.

These lists contain the UniProt identifiers for each of the heteromeric ligands and receptors (“uniprot1”, “uniprot2”,...) and a name given to the complex (“complex_name”). Common fields with “proteins_input” include: “transmembrane”, “peripheral”, “secreted”, “secreted_desc”, “secreted_highlight”, “receptor”, “receptor_desc”, “integrin”, “other”, “other_desc” (see description in the above “protein_input” section for clarification). In addition, we include other optional information that may be relevant for the stoichiometry of the heterodimers. If heteromers are defined in the RCSB Protein Data Bank (http://www.rcsb.org/), structural information is included in our CellPhoneDB annotation in the “pdb_structure”, “pdb_id” and “stoichiometry”. An additional field “comments_complex” was created to add a brief description of the heteromer.

#### 4. “interaction_input”

*Mandatory fields: “partner_a”; “partner_b”; “annotation_strategy”; “source”*

*Optional fields: “protein_name_a”; “protein_name_b”*

Interactions stored in CellPhoneDB are annotated using their UniProt identifier (binary interactions) or the name of the complex (interactions involving heteromers) (“partner_a” and “partner_b”). The name of the protein is also included, yet not mandatory (“protein_name_a” and “protein_name_b”). Protein names are not stored in the database.

There are two main inputs of interactions: i) a systematic input querying other databases, and ii) a manual input using curated information from CellPhoneDB developers (“interactions_curated”) and users. The method used to assign the interaction is indicated in the “annotation_strategy” column.

Each interaction stored has a CellPhoneDB unique identifier (“id_cp_interaction”) generated automatically by the internal pipeline.

##### a) Systematic input from other databases

We consider interacting partners as: (i) binary interactions annotated by IUPHAR (http://www.guidetopharmacology.org/) and (ii) cytokines, hormones and growth factors interactions annotated by innateDB (https://www.innatedb.com/) and the iMEX consortium (https://www.imexconsortium.org/).

Binary interactions from IUPHAR are directly downloaded from “http://www.guidetopharmacology.org/DATA/interactions.csv” and “guidetopharmachology.org” is indicated in the “annotation_strategy” field. For the iMEX consortium all protein-protein interactions are downloaded using the PSICQUIC REST APIs^19^. The IMEx^20^, IntAct ^21^, InnateDB ^22^, UCL-BHF, MatrixDB ^23^, MINT ^24^, I2D ^25^, UniProt, MBInfo registries are used. Interacting partners are defined as follows:

- Interacting partner A has to be transmembrane, receptor and cannot be classified as “others” (see the previous “protein_list” section for more information).
- Interacting partner B has to be “secreted_highlight”. This group of proteins includes cytokines, hormones, growth factors and other immune-related proteins (see the previous “protein_list” section for more information).

In both cases, some interactions are excluded: i) interactions where one of the components is part of a complex (see “complex_curated” list in the above section); ii) interactions which are not involved in cell-cell communication or are wrongly annotated by our systematic method. These are stored in a curated list of proteins named “excluded_interaction”. The “excluded_interaction” file contains five fields: a) uniprot_1: name of the interacting partner A that is going to be excluded; b) uniprot_2: name of the interacting partner B that is going to be excluded; c) name.1: name of the protein to be excluded corresponding to uniprot_1; d) name.2: name of the protein to be excluded corresponding to uniprot_2; e) comments: information about the exclusion of the protein.

Homomeric complexes - proteins interacting with themselves - are excluded from the systematic analysis. Importantly, in cases where both the systematic and the curated input detect the interactions, the curated input always prevails over the systematic information.

##### b) Curated approach

The majority of ligand–receptor interactions are manually curated by reviewing UniProt descriptions and PubMed information on membrane receptors. Cytokine and chemokine interactions were annotated following the International Union of Pharmacology annotation ^26^. The interactions of other groups of cell-surface proteins were manually reviewed, including the TGF family, integrins, lymphocyte receptors, semaphorins, ephrins, Notch and TNF receptors. The bibliography used to annotate the interaction is stored in “source”. ‘Uniprot’ indicates that the interaction has been annotated using UniProt descriptions.

#### User-defined receptor-ligand datasets

Our system allows users to create their own lists of curated proteins and complexes. In order to do so, the format of the users’ lists must be compatible with the input files. Users can submit their lists using the Python package version of CellPhoneDB, and then send them via email, the cellphonedb.org form, or a pull request to the CellPhoneDB data repository (https://github.com/Teichlab/cellphonedb-data).

### Database structure

Information is stored in an SQLite relational database (https://www.sqlite.org/). SQLAlchemy (www.sqlalchemy.org) and Python 3 were used to build the database structure and the query logic. The application is designed to allow analysis on potentially large count matrices to be performed in parallel. This requires an efficient database design, including optimisation for query times, indices and related strategies. All application code is open source and uploaded both to github and the web server. An explanation of the content of the tables and the database schema is available in Supplementary methods and Supplementary Figure 1 and 2.

### Statistical inference of receptor-ligand specificity

To assess cellular crosstalk between different cell types, we use our repository in a statistical framework for inferring cell–cell communication networks from single-cell transcriptome data. We predict enriched receptor–ligand interactions between two cell types based on expression of a receptor by one cell type and a ligand by another cell type, using scRNA-seq data. To identify the most relevant interactions between cell types, we look for the cell-type specific interactions between ligands and receptors. Only receptors and ligands expressed in more than a user-specified threshold percentage of the cells in the specific cluster are considered significant (default is 10%).

We then perform pairwise comparisons between all cell types. First, we randomly permute the cluster labels of all cells (1,000 times as a default) and determine the mean of the average receptor expression level in a cluster and the average ligand expression level in the interacting cluster. For each receptor–ligand pair in each pairwise comparison between two cell types, this generates a null distribution. By calculating the proportion of the means which are as or higher than the actual mean, we obtain a *p*-value for the likelihood of cell-type specificity of a given receptor–ligand complex. We then prioritize interactions that are highly enriched between cell types based on the number of significant pairs, so that the user can manually select biologically relevant ones. For the multi-subunit heteromeric complexes, we require that all subunits of the complex are expressed (using a user-specified threshold), and therefore we use the member of the complex with the minimum average expression to perform the random shuffling.

### Cell subsampling for accelerating analyses

Technological developments and protocol improvements have enabled an exponential growth of the number of cells obtained from scRNA-seq experiments^27^. Large-scale datasets can profile hundreds of thousands cells, which presents a challenge for the existing analysis methods in terms of both memory usage and runtime. In order to improve the speed and efficiency of our protocol and facilitate its broad accessibility, we integrated subsampling as described in Hie et al.^28^. This “geometric sketching” approach aims to maintain the transcriptomic heterogeneity within a dataset with a smaller subset of cells. The subsampling step is optional, enabling users to perform the analysis either on all cells, or with other subsampling methods of their choice.

## Procedure

### Input data files and software required

- META file: Cell type annotation. Contains two columns: “Cell” indicating the name of the cell, and “cell_type” indicating the name of the cluster considered. Format accepted: .csv, .txt, .tsv, .tab, pickle.
- COUNTS file: scRNA-seq count data containing gene expression values where rows are genes presented with gene names identifiers (Ensembl IDs, gene names or hgnc_symbol annotation) and columns are cells. We recommend using normalised count data. Importantly, when using the subsampling option, the user needs to specify whether the data was log-transformed. Format accepted: .csv or .txt, .tsv, .tab, pickle.

Example input data can be downloaded from our webserver at https://www.cellphonedb.org/explore-sc-rna-seq or by running:

curl https://raw.githubusercontent.com/Teichlab/cellphonedb/master/in/example_data/test_counts.txt--outputtest_counts.txt

curl https://raw.githubusercontent.com/Teichlab/cellphonedb/master/in/example_data/test_meta.txt--outputtest_meta.txt

#### Software

Python 3.5 or higher, SQLAlchemy, SQLite.

#### Pre-processing of raw data and input files for the protocol

We recommend using normalised count data. This can be obtained by taking the raw data from the Seurat object and applying the normalisation manually. The user can also normalise using their preferred method.

~~~
*# take raw data and normalise it*
count_raw <-data_object@raw.data[,data_object@cell.names]
count_norm <-apply(count_raw, 2, function(x) (x/sum(x))*10000)
write.table(count_norm, ‘cellphonedb_count.txt’, sep=’\t’, quote=F)
*# generating meta file*
meta_data <-cbind(rownames(data_object@meta.data), data_object@meta.data[,’cluster’, drop=F]) *# cluster is the user’s corresponding cluster column*
write.table(meta_data, ‘cellphonedb_meta.txt’, sep=’\t’, quote=F, row.names=F)
~~~

The input files can also be extracted from a scanpy adata object:

~~~
import pandas as pd
import scanpy.api as sc
*# data after filtering and normalising*
adata = sc.read(adata_filepath)
# *we recommend using the normalised non-log transformed data - you can save it in adata.norm for example*
df_expr_matrix = adata.norm
df_expr_matrix = df_expr_matrix.T
df_expr_matrix = pd.DataFrame(df_expr_matrix.toarray())
*# Set cell ids as columns*
df_expr_matrix.columns = adata.obs.index
*# Genes should be either Ensembl IDs or gene names*
df_expr_matrix.set_index(adata.raw.var.index, inplace=True)
df_expr_matrix.to_csv(savepath_counts,sep=’\t’)
*# generating meta file*
df_meta = pd.DataFrame(data={’Cell’: list(adata.obs[cell_ids]), ‘cell_type’: list(adata.obs[annotation_name])})
df_meta.set_index(’Cell’,inplace=True)
df_meta.to_csv(savepath_meta, sep=’\t’)
~~~

CellPhoneDB can be used either through the interactive website (cellphonedb.org) which executes calculations in our private cloud, or as a Python package using the user’s computer/cloud/farm. The Python package is recommended for big datasets.

## Python package

### Installation

Note: If the default Python interpreter is for v2.x (can be checked with the command: python --version), calls to python/pip must be substituted by python3/pip3.

We highly recommend using a virtual environment (steps 1 and 2), but can be omitted.

1. Create python > 3.5 virtual-env python -m venv cpdb-venv
2. Activate virtual-env source cpdb-venv/bin/activate
3. Install cellphonedb pip install cellphonedb

### Running with statistical analysis

4. Activate virtual-env if you have not activated previously source cpdb-venv/bin/activate
5. Run in statistical analysis mode using the input files for metadata and counts cellphonedb method statistical_analysis test_meta.txt test_counts.txt

Optional parameters:

--project-name: Name of the project. It creates a subfolder in output folder

--iterations: Number of pvalues analysis iterations [1000]

--threshold: % of cells expressing a gene

--result-precision: Number of decimal digits in results [3]

--output-path: Directory where the results will be allocated (the directory must exist) [out]

--means-result-name: Name of the means result file [means.txt]

--significant-mean-result-name: Name of the significant means result file [significant_means.txt]

--deconvoluted-result-name: Name of the deconvoluted result file [deconvoluted.txt]

--verbose/--quiet: Print or hide cellphonedb logs [verbose]

--pvalues-result-name: Name of the pvalues result file [pvalues.txt]

--debug-seed: Debug random seed −1 for disable it. >=0 [-1]

--threads: Number of threads to use. >=1 [-1]

### Usage Examples

Set number of iterations and threads:

cellphonedb method statistical_analysis yourmetafile.txt yourcountsfile.txt --iterations=10 -- threads=2

Set project subfolder:

cellphonedb method analysis yourmetafile.txt yourcountsfile.txt --project-name=new_project

Set output path:

mkdir custom_folder

cellphonedb method statistical_analysis yourmetafile.txt yourcountsfile.txt --output-path=custom_folder

### Running with subsampling and statistical analysis

6. Run in statistical analysis mode using the input files for metadata and counts and add --subsampling + other subsampling specific parameters

~~~
cellphonedb method statistical_analysis yourmetafile.txt yourcountsfile.txt -- subsampling --subsampling-log true
Subsampling specific parameters:
--subsampling-log: Enable log transformation for non log transformed data inputs (mandatory parameter)
--subsampling-num-pc: Subsampling NumPC argument
--subsampling-num-cells: Number of cells to subsample (defaults to a 1/3 of cells)
~~~

### Running without statistical analysis

7. Run in normal mode using the input files for metadata and counts and specified -- threshold parameter. The parameters are as described above, with the exception of --

~~~
pvalues-result-name and --debug-seed
cellphonedb method analysis test_meta.txt test_counts.txt
~~~

### Visualisation

8. Run after running cellphonedb in either statistical analysis mode or normal mode using the means.csv and pvalues.scv output files

~~~
cellphonedb plot dot_plot
Dot plot specific parameters:
--means-path: Analysis output means [./out/means.txt]
--pvalues-path: Analysis output pvalues [./out/pvalues.txt]
--output-path: Output folder [./out]
--output-name: Name of the output plot [plot.pdf]; available output formats are those supported by R’s ggplot2 package, among others are: pdf, png, jpeg
--rows: File with a list of rows to plot, one per line [all available]
--columns: File with a list of columns to plot, one per line [all available]
--verbose / --quiet: Print or hide cellphonedb logs [verbose]
~~~ To plot only desired rows/columns: cellphonedb plot dot_plot --rows in/rows.txt --columns in/columns.txt
9. Run after running cellphonedb in either statistical analysis mode or normal mode using the the pvalues.scv output file. cellphonedb plot heatmap_plot meta_data Heatmap plot specific parameters:

~~~
--pvalues-path: Analysis output pvalues [./out/pvalues.txt]
--output-path: Output folder [./out]
--count-name: Name of the output plot [heatmap_count.pdf]
--log-name: Name of the output log plot [heatmap_log_count.pdf]
--verbose / --quiet: Print or hide cellphonedb logs [verbose]
~~~
10. The user can run the analysis using a user-specified version of the database cellphonedb method statistical_analysis in/example_data/test_meta.txt in/example_data/test_counts.txt --database=v0.0.2
11. Detailed description of the mandatory and optional parameters can be found using the help option: cellphonedb method statistical_analysis yourmetafile.txt yourcountsfile.txt --help

## Interactive web portal

The web interface includes form inputs for the user to decide on analysis parameters before submission. Downstream calculations are performed on the application’s servers, rendering the information of ligand and receptor expression, and visualisation diagrams once analysis is complete (Figure 5).

**Figure 5.**
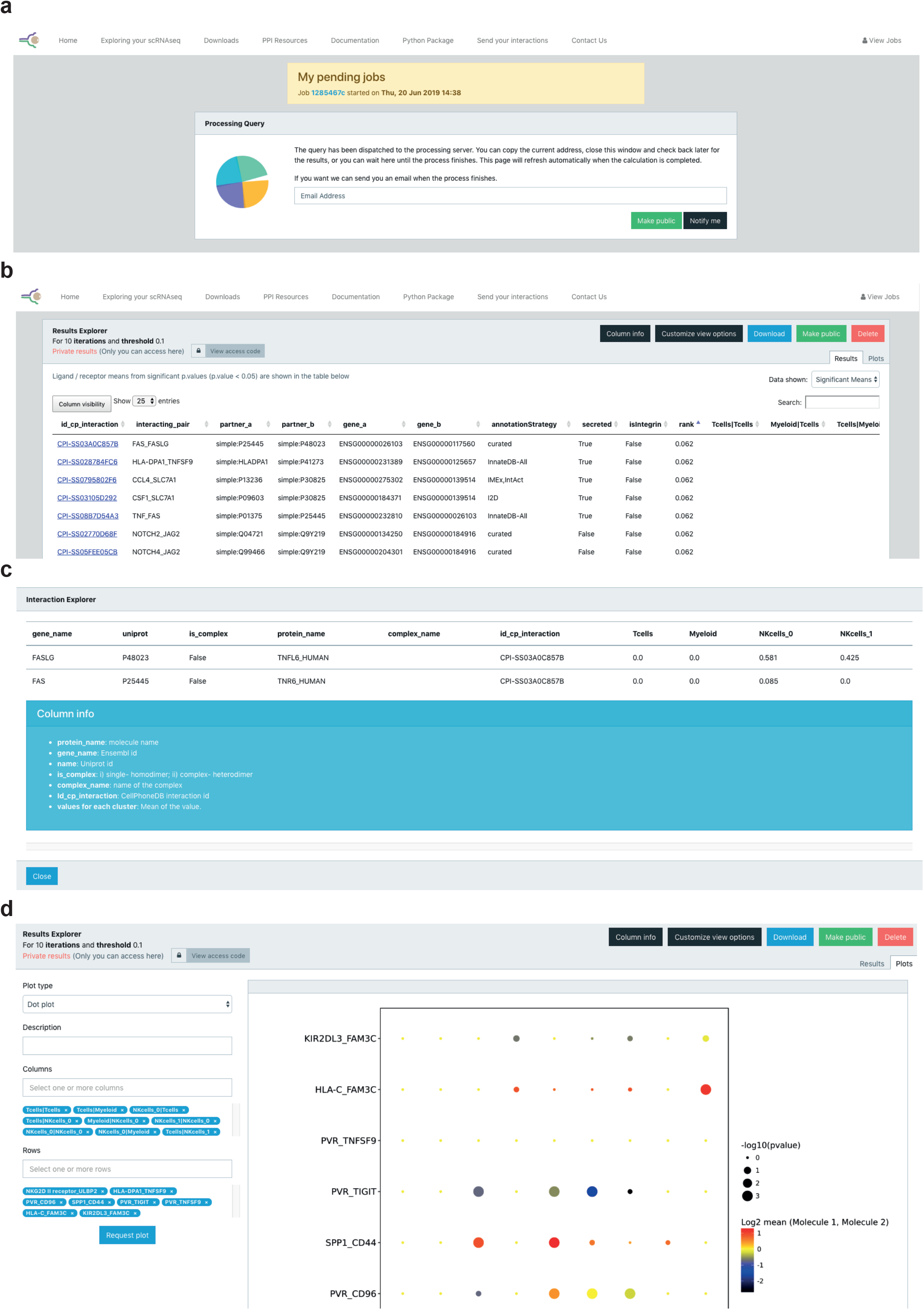
Screenshot of web portal. **a**, Screenshot showing how to input the email in order to get notification when the job is done. **b**, Screenshot showing the significant_means results table. The user can click on a selected id_cp_interaction field to get a more detailed information for the specific interaction pair. **c**, Screenshot showing detailed information for the specific interaction pair that appears when the user clicks on a specific id_cp_interaction field. **d**, Screenshot showing the dot plot visualisation page.

12. Go to the tab “Exploring your scRNAseq” and input the meta and count input files.
13. Provide an email address if you would like to get an update when the process finishes (Figure 5a).
14. The significant means results table will appear as in Figure 5c. You can change the current view by clicking on the:Data Shown” button (Figure 5b) and can download the results as well. By clicking on any field from the id_cp_interaction column, detailed information for the specific interaction pair will appear (Figure 5c).
15. Go to the tab “Plots” and pick the type of plot you would like to produce. For plotting dot plots, please select the columns and rows you need (Figure 5d).

The online results viewer allows you to select which columns you wish to display in each table. This option is quite useful as an aid to visualize the results.

## Anticipated results: formats of files

We originally applied CellPhoneDB to study the maternal-fetal communication at the decidual-placental interface during early pregnancy^1^. The results obtained with our new CellPhoneDB v2.0 using subsampling were consistent with our original conclusions (Figure 3). Here we provide an explanation of the results generated in this example.

Without running statistical inference of receptor-ligand interactions, only “means.csv” and “desconvoluted.csv” are generated. The “means.csv” file contains mean values for each ligand-receptor interaction. The “deconvoluted.csv” file gives additional information for each of the interacting partners. This is important as some of the interacting partners are heteromers. In other words, multiple molecules have to be expressed in the same cluster in order for the interacting partner to be functional. If the user uses the statistical inference approach, additional “pvalues.csv” and “significant_means.csv” file are generated with the values for the significant interactions.

It is important to bear in mind that the interactions are not symmetric. In other words, when testing a ligand/receptor pair A_B between clusters X_Y, the expression of partner A is considered within the first cluster (X), and the expression of partner B within the second cluster (Y). Therefore, X_Y and Y_X represent different comparisons and will have different *p*-values and means.

### Code availability

CellPhoneDB code is available at https://github.com/Teichlab/cellphonedb. It can also be downloaded from https://cellphonedb.org/downloads.

### Data availability

The decidua and placenta datasets were deposited in ArrayExpress, with experiment code E-MTAB-6701.

## Acknowledgments

We thank Kerstin Meyer and Mike Stubbington for scientific discussions, Pablo Porras for advice on querying the IMEx database, Luz Garcia-Alonso and Krzysztof Polanski for carefully reading the manuscript, Gavin J Wright, Laura Wood and Gerard Graham for advice on protein-protein interactions and Jana Eliasova and Ania Hupalowska for help with the illustrations. We are grateful to Adria Lopez and YDEVS members for their help with the webserver and the implementation of the code in github, and all the Teichmann lab and Vento-Tormo lab members for their fruitful advice. The project was supported by Wellcome Sanger core funding (no. WT206194).

## Author contribution

M.E, S.A.T and R.V-T conceived and developed the protocol and wrote the manuscript. M.V-T developed the protocol, implemented the code in the web server and github and contributed to writing the manuscript.

---

**Table 1.**
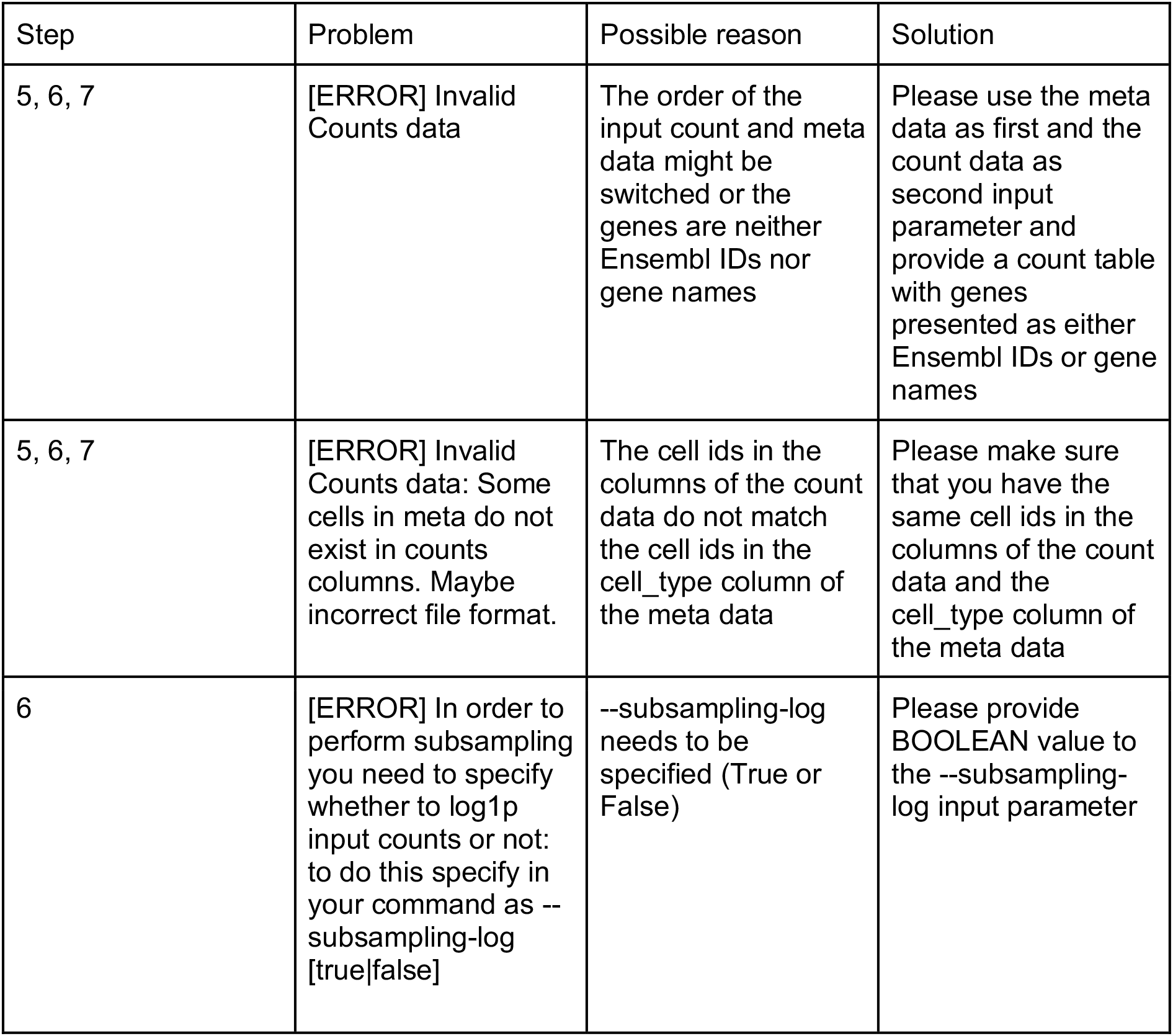
Troubleshooting table.

**Table 2.**
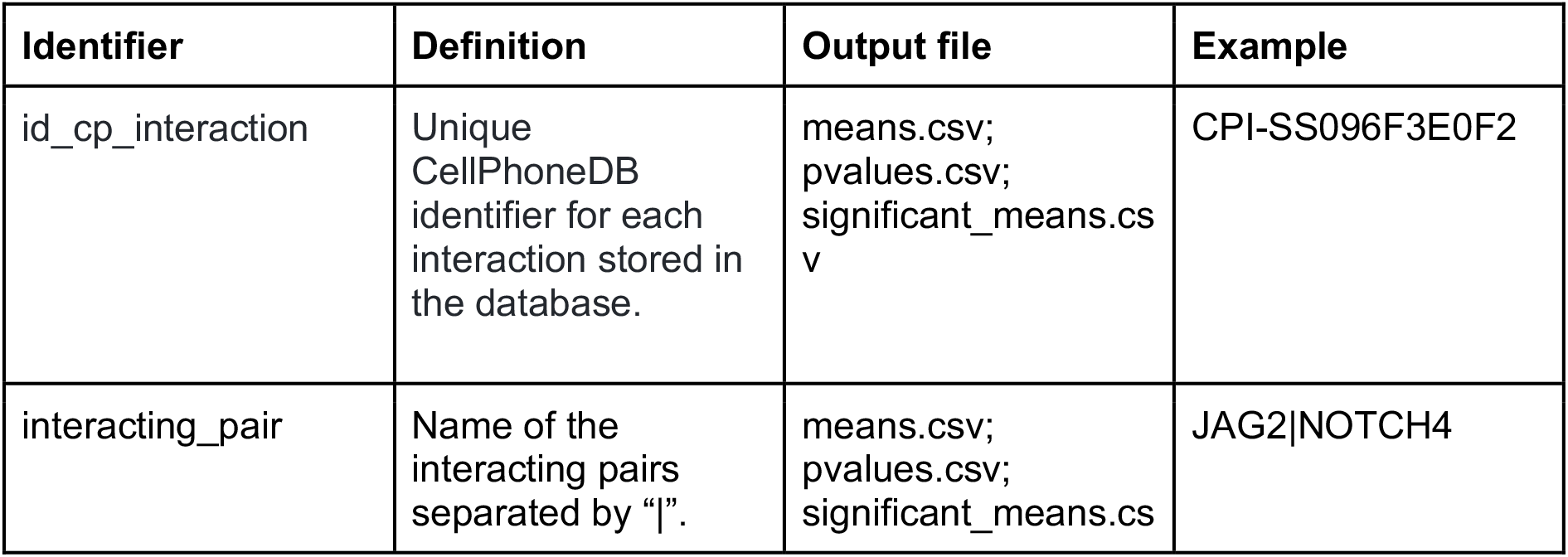
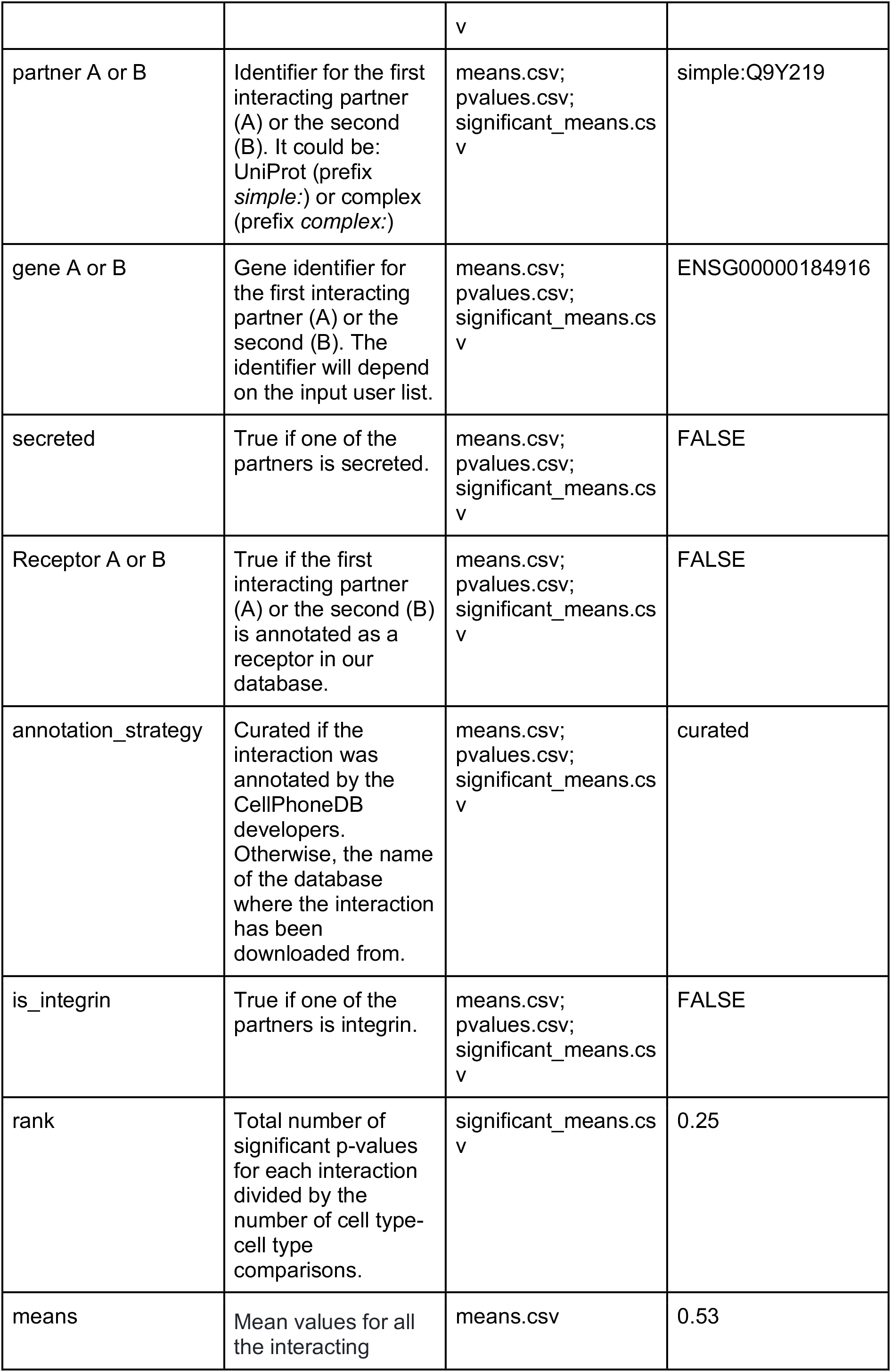
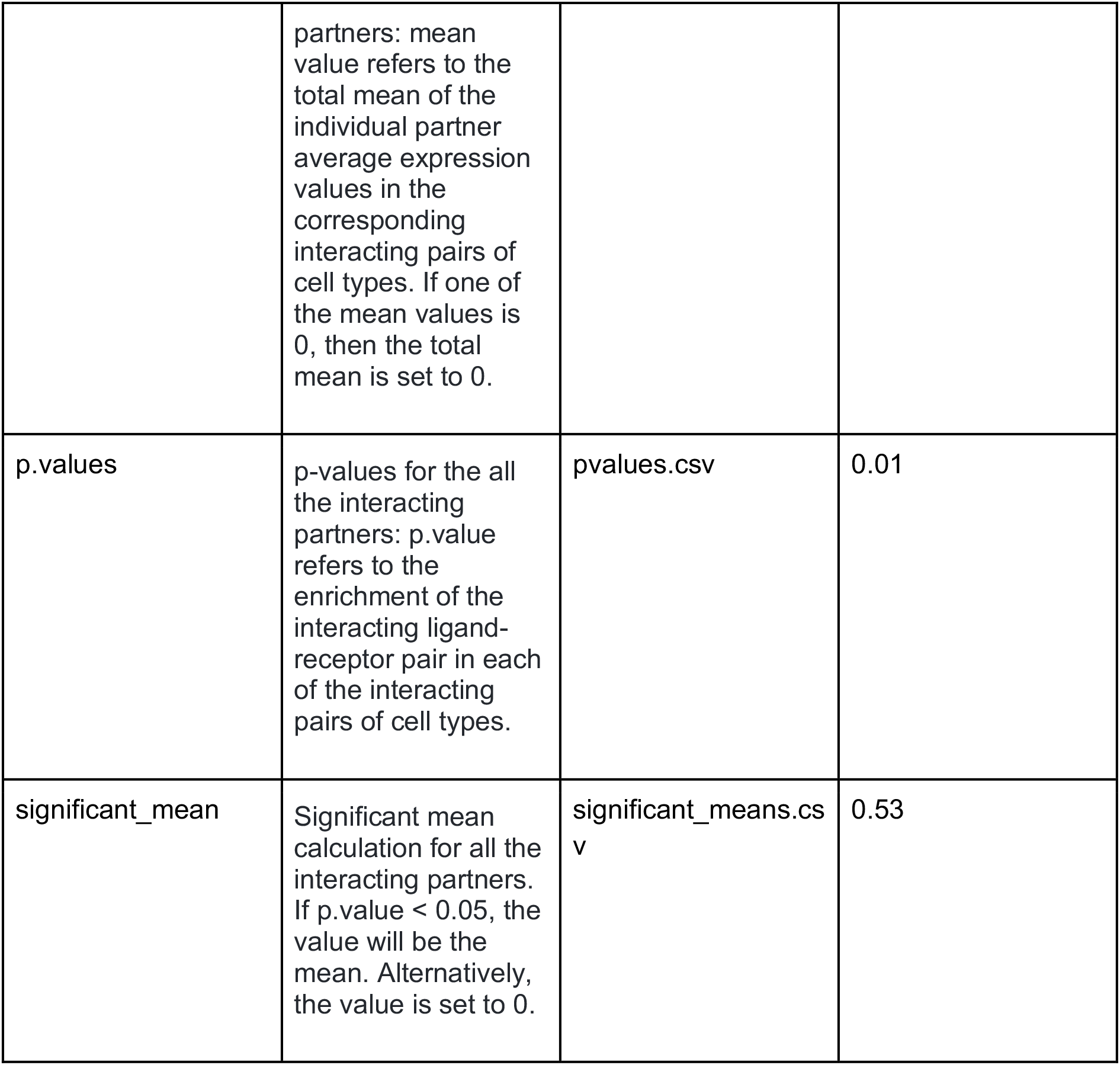
Description of the output files means.csv, pvalues.csv and significant_means.csv.

**Table 3.**
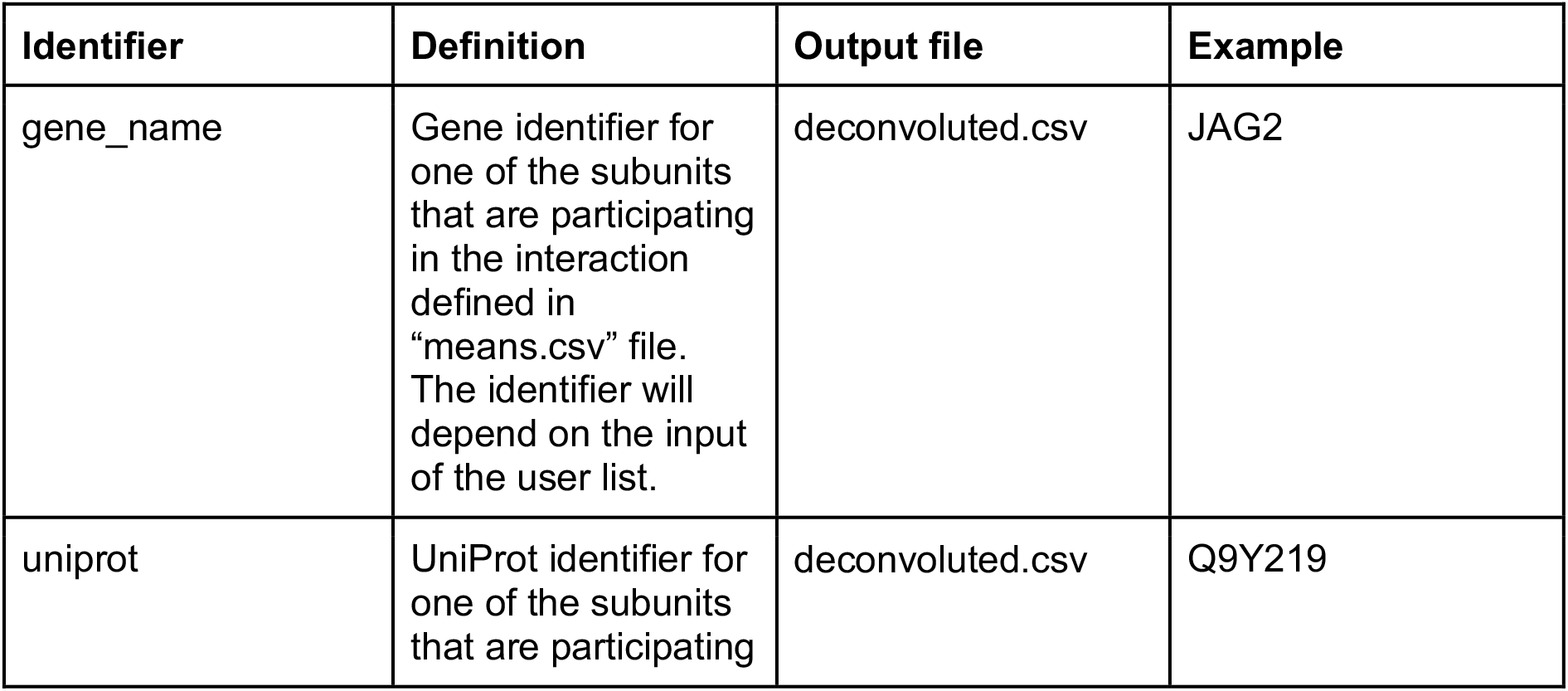
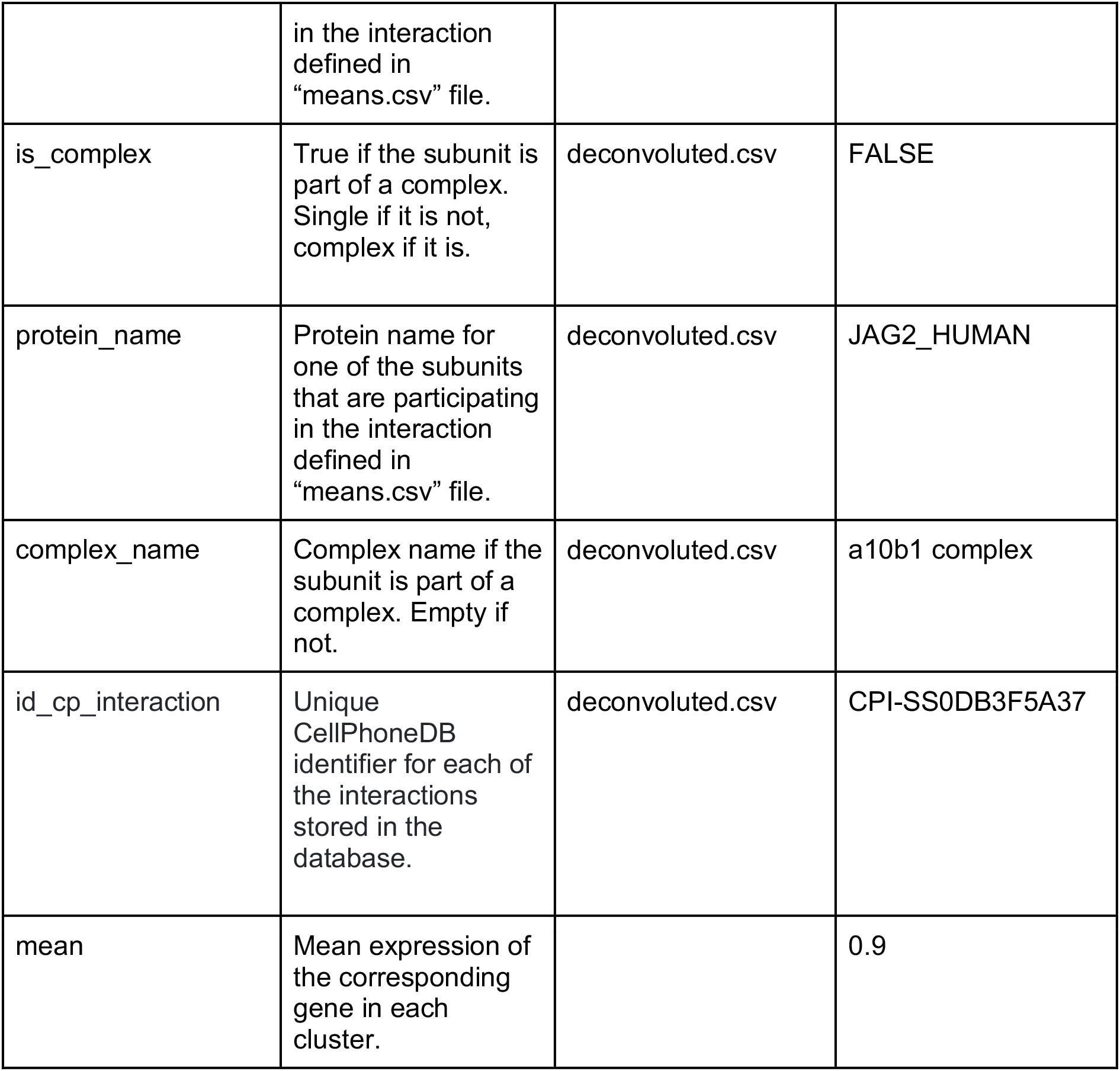
Description of the output file deconvoluted.csv.

| Identifier | Definition | Output file | Example |
| --- | --- | --- | --- |
| gene_name | Gene identifier for one of the subunits that are participating in the interaction defined in "means.csv" file. The identifier will depend on the input of the user list. | deconvoluted.csv | JAG2 |
| uniprot | UniProt identifier for one of the subunits that are participating in the interaction defined in "means.csv" file. | deconvoluted.csv | Q9Y219 |
| is_complex | True if the subunit is part of a complex. Single if it is not, complex if it is. | deconvoluted.csv | FALSE |
| protein_name | Protein name for one of the subunits that are participating in the interaction defined in "means.csv" file. | deconvoluted.csv | JAG2_HUMAN |
| complex_name | Complex name if the subunit is part of a complex. Empty if not. | deconvoluted.csv | a10b1 complex |
| id_cp_interaction | Unique CellPhoneDB identifier for each of the interactions stored in the database. | deconvoluted.csv | CPI-SS0DB3F5A37 |
| mean | Mean expression of the corresponding gene in each cluster. |  | 0.9 |

## Supplementary information

### Supplementary methods

The database consists of 6 main tables: gene_table; protein_table; multidata_table; interaction_table; complex_table; complex_composition_table (Supplementary Figure 1).

All tables have an incremental numeric unique identifier with the structure id_{table_name} and one or more foreign keys, with structure {foreign_table_name}_id, to connect all tables.

#### 1. gene_table

This table stores all the information generated in the gene_input database input file. This includes the gene name (“gene_name”); the HUGO nomenclature committee symbol (HGNC) (“hgnc_symbol”) and ensembl identifier (“ensembl”). Importantly, only the gene and protein information of proteins considered in the “interactions_list” is stored in our database.

The gene table is related to the protein table via the protein_id - id_protein (one to many) foreign key.

#### 2. multidata_table

This table stores the shared information between the protein_table and the complex_table and represents (in terms of database design and performance) the relations in the interaction_table.

All the information required in this table is obtained from the *protein_input* and *complex_input* input files. It stores the following fields: *name* (it corresponds to *uniprot* if the row is a protein or *complex_name* if row is a complex), *transmembrane*, *peripheral*, *secreted*, *secreted_desc*, *secreted_highlight*, *receptor*, *receptor_desc*, *integrin*, *other* and *other_desc*. *is_complex* column was added for internal optimization and indicates if the row is a complex.

#### 3. protein_table

This table stores the information obtained from the database input file *protein_input*. It contains the name of the protein (*protein_name*), *tags*, *tags_reason* and *tags_description*. The table is related to *multidata_table* (1..0 - 1 relation) through the *protein_multidata_id* foreign key.

#### 4. complex_table

This table stores complex information from the database input file *complex_input*. It stores the following fields: *pdb_id*, *pdb_structure*, *stoichiometry*, *comments_complex*. The table is related to *multidata_table* (1..0 - 1 relation) through the *complex_multidata_id* foreign key.

All information about the complex components is stored in the *complex_composition_table*.

#### 5. complex_composition_table

This table stores the proteins that compose (*uniprot_1* - *uniprot_4*) a complex. It is related to multidata through *complex_multidata_id* and to *multidata_table* through *protein_multidata_id* (* to 1 relations). We also created an additional column called *total_protein* (with number of complex components) for internal optimization purposes. Supplementary Figure 2 represents an example of two *complex_input* rows with two and four components.

#### 6. interaction_table

This table stores the interactions data from *interaction_input* file. It uses the following columns to represent the data: *id_cp_interaction*, *annotation_strategy* and *source*. To identify the interaction partners (*partner_a* and *partner_b* in *interaction_input*), the table is connected to *multidata_table* through the foreign key *multidata_1_id* and *multidata_2_id* respectively with 1 - 1..* relation. Since *multidata_table* stores both protein and complex data, it is not necessary to create additional columns to show if components *partner_a* and *partner_b* are simple or complex. Importantly, only genes and proteins involved in cell-cell communication are stored in our database (see the *interaction_list* section).

## Supplementary figures

**Supplementary Figure 1.**
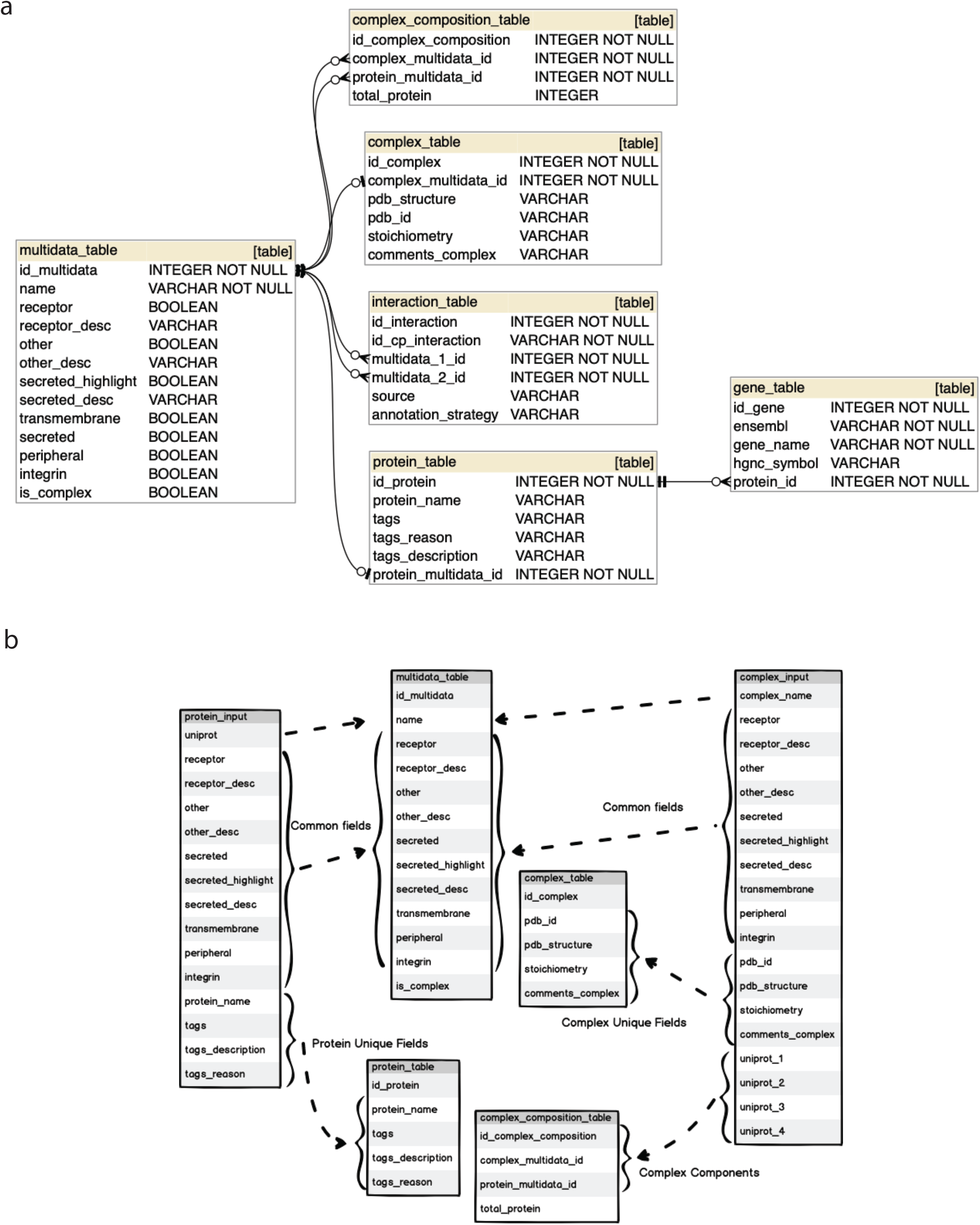
Diagram database structure. **a)** database schema, **b)** protein_input/complex_input storage in the CellPhoneDB database tables. The multidata entity stores fields common to complex_input and protein_input. This makes it easier and faster for the user to perform interaction queries because interaction_table is only related to multidata_table. There is no need to check if the component is simple or complex to find the correct table. All non-common fields are stored in either protein_table or complex_table. Complex fields are stored in complex_composition_table as it’s typical in relational databases. The is_complex and total_protein field are created for optimization purposes.

**Supplementary Figure 2.**
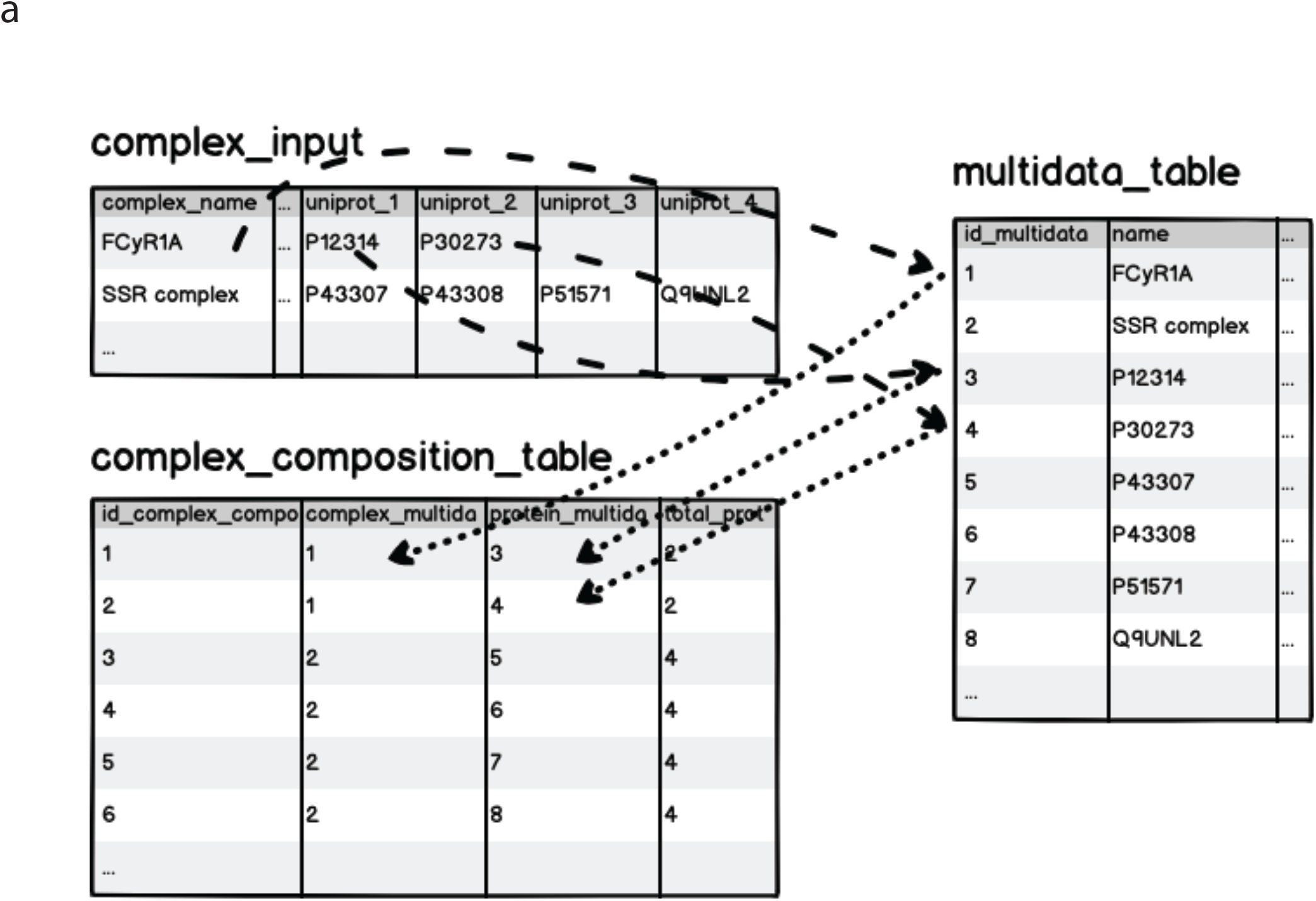
Example of complex_input components stored in CellPhoneDB. **a**, An example of two *complex_input* rows with two and four components.

## References

1. Vento-Tormo, R. et al. Single-cell reconstruction of the early maternal-fetal interface in humans. Nature 563, 347–353 (2018).

2. Braga, F. A. V. et al. A cellular census of human lungs identifies novel cell states in health and in asthma. Nature Medicine (2019). doi: 10.1038/s41591-019-0468-5

3. Davidson, S. et al. Single-cell RNA sequencing reveals a dynamic stromal niche within the evolving tumour microenvironment. doi: 10.1101/467225

4. Lubeck, E., Coskun, A. F., Zhiyentayev, T., Ahmad, M. & Cai L. Single-cell in situ RNA profiling by sequential hybridization. Nat. Methods 11, 360–361 (2014).

5. Ståhl, P. L. et al. Visualization and analysis of gene expression in tissue sections by spatial transcriptomics. Science 353, 78–82 (2016).

6. Rodriques, S. G. et al. Slide-seq: A scalable technology for measuring genome-wide expression at high spatial resolution. Science 363, 1463–1467 (2019).

7. Svensson, V. A method for transcriptome-wide gene expression quantification in intact tissues. Immunol. Cell Biol. 97, 439–441 (2019).

8. Skelly, D. A. et al. Single-Cell Transcriptional Profiling Reveals Cellular Diversity and Intercommunication in the Mouse Heart. Cell Rep. 22, 600–610 (2018).

9. Camp, J. G. et al. Multilineage communication regulates human liver bud development from pluripotency. Nature 546, 533–538 (2017).

10. Pavličev, M. et al. Single-cell transcriptomics of the human placenta: inferring the cell communication network of the maternal-fetal interface. Genome Res. 27, 349–361 (2017).

11. Puram, S. V. et al. Single-Cell Transcriptomic Analysis of Primary and Metastatic Tumor Ecosystems in Head and Neck Cancer. Cell 171, 1611–1624.e24 (2017).

12. Suryawanshi, H. et al. A single-cell survey of the human first-trimester placenta and decidua. Sci Adv 4, eaau4788 (2018).

13. Zhou, J. X., Taramelli, R., Pedrini, E., Knijnenburg, T. & Huang, S. Author Correction: Extracting Intercellular Signaling Network of Cancer Tissues using Ligand-Receptor Expression Patterns from Whole-tumor and Single-cell Transcriptomes. Sci. Rep. 8, 17903 (2018).

14. Cohen, M. et al. Lung Single-Cell Signaling Interaction Map Reveals Basophil Role in Macrophage Imprinting. Cell 175, 1031–1044.e18 (2018).

15. Kumar, M. P. et al. Analysis of Single-Cell RNA-Seq Identifies Cell-Cell Communication Associated with Tumor Characteristics. Cell Rep. 25, 1458–1468.e4 (2018).

16. Halpern, K. B. et al. Paired-cell sequencing enables spatial gene expression mapping of liver endothelial cells. Nat. Biotechnol. 36, 962–970 (2018).

17. Joost, S. et al. Single-Cell Transcriptomics of Traced Epidermal and Hair Follicle Stem Cells Reveals Rapid Adaptations during Wound Healing. Cell Rep. 25, 585–597.e7 (2018).

18. Boisset, J.-C. et al. Mapping the physical network of cellular interactions. Nat. Methods 15, 547–553 (2018).

19. Proteomics Standards Initiative Common Query Interface. Encyclopedia of Systems Biology 1798–1798 (2013). doi: 10.1007/978-1-4419-9863-7_101243

20. Orchard, S. et al. Protein interaction data curation: the International Molecular Exchange (IMEx) consortium. Nat. Methods 9, 345–350 (2012).

21. Orchard, S. et al. The MIntAct project--IntAct as a common curation platform for 11 molecular interaction databases. Nucleic Acids Res. 42, D358–63 (2014).

22. Breuer, K. et al. InnateDB: systems biology of innate immunity and beyond--recent updates and continuing curation. Nucleic Acids Res. 41, D1228–33 (2013).

23. Clerc, O. et al. MatrixDB: integration of new data with a focus on glycosaminoglycan interactions. Nucleic Acids Res. 47, D376–D381 (2019).

24. Licata, L. et al. MINT, the molecular interaction database: 2012 update. Nucleic Acids Res. 40, D857–61 (2012).

25. Brown, K. R. & Jurisica, I. Unequal evolutionary conservation of human protein interactions in interologous networks. Genome Biol. 8, R95 (2007).

26. Bachelerie, F. et al. International Union of Basic and Clinical Pharmacology. [corrected]. LXXXIX. Update on the extended family of chemokine receptors and introducing a new nomenclature for atypical chemokine receptors. Pharmacol. Rev. 66, 1–79 (2014).

27. Svensson, V., Vento-Tormo, R. & Teichmann, S. A. Exponential scaling of single-cell RNA-seq in the past decade. Nat. Protoc. 13, 599–604 (2018).

28. Hie, B., Cho, H., DeMeo, B., Bryson, B. & Berger, B. Geometric Sketching Compactly Summarizes the Single-Cell Transcriptomic Landscape: Supplementary Information. doi: 10.1101/536730

